# Development of diagnostic PCR and LAMP markers for *MALE STERILITY 1* (*MS1*) in *Cryptomeria japonica* D. Don

**DOI:** 10.1101/2020.05.19.090092

**Authors:** Yoichi Hasegawa, Saneyoshi Ueno, Fu-Jin Wei, Asako Matsumoto, Tokuko Ujino-Ihara, Kentaro Uchiyama, Yoshinari Moriguchi, Masahiro Kasahara, Takeshi Fujino, Shuji Shigenobu, Katsushi Yamaguchi, Takahiro Bino, Tetsuji Hakamata

## Abstract

**Objective:** Due to the allergic nature of the pollen of *Cryptomeria japonica*, the most important Japanese forestry conifer, a pollen-free cultivar is preferred. Mutant trees detected in nature have been used for the production of a pollen-free cultivar. In order to reduce the time and cost needed for the production and breeding, we aimed to develop simple diagnostic molecular markers for mutant alleles of the causative gene *MALE STERILITY 1* (*MS1*) in *C. japonica*. The expected function of this gene, its two dysfunctional mutations, and genetic diversity were described recently in a related study.

**Results:** We have developed PCR and LAMP markers to detect mutant alleles and to present experimental options depending on available laboratory equipment. At field stations, where PCR machines are not available, LAMP markers were developed. LAMP only needs heat-blocks or a water bath to perform the isothermal amplification and assay results can be easily seen by eye. Because the causative mutations were deletions, two kinds of PCR markers, amplified length polymorphism (ALP) and allele specific PCR (ASP) markers, were developed. These assays can be carried out by capillary or agarose gel electrophoresis.

## Introduction

Molecular markers allow for the selection of specific phenotypes once the associated genotype and candidate gene(s) are identified. Due to the large body mass of tree species, especially conifers, maker-assisted selection with the candidate gene at early life stage (seedling) is helpful in reducing the cost and time of breeding. Sugi (*Cryptomeria japonica* D. Don) is the most important conifer in Japan, occupying about 40% of artificial forests. Due to its fast growth and straight bowl, sugi timber has been used not only for building materials, but also for daily life consumables such as chopsticks and bowls. Recently, however, the pollen of sugi has caused pollenosis, which affects approximately 25% of Japanese citizens [1]. Therefore, breeding efforts are currently focused on producing a sterile pollen cultivar [2]. A candidate gene for male sterility, *MALE STERILITY 1* (*MS1*), has been identified by transcriptome analysis and linkage mapping. Diagnostic molecular markers are useful in laboratories as well as in forest stations in search for the mutant alleles (*ms1*) from the vast amount of genetic resources. Such markers would be useful in breeding for male sterile sugi.

Here, we developed diagnostic markers for *MS1* in sugi for use in laboratories as well as in the forest stations. Amplicon length polymorphism (ALP) and allele specific PCR (ASP) markers were developed for laboratories equipped with capillary and/or gel electrophoretic systems. These markers are suitable for precise determination of genotypes. For nurseries and forest stations, we developed LAMP (loop-mediated isothermal amplification) [3] primer sets. LAMP experiments are less resource intensive and enable the rapid detection of alleles in the field without the need for a molecular biology laboratory. Utilization of these markers in breeding projects for male sterility will boost the production of male sterile cultivars, and hopefully reduce the allergic pollen cloud in the future.

## Main text

## Materials and Methods

Three individual trees with known genotypes for *MS1*: ‘Fukushima1’ (*ms1-1*/*ms1-1*), ‘Ooi-7’ (*Ms1*/*ms1-2*), and ‘G492’ (*Ms1*/*ms1-1*) were used from a mapping family of *MS1* [4]. ‘G492’ is an offspring of the cross between ‘Fukushima1’ (male sterile seed parent) and ‘Ooi-7’ (male fertile pollen parent). For ‘Fukushima1’ and ‘Ooi-7’, sequencing analysis had been performed for *MS1* gene (CJt020762), confirming a 4-base pair (bp) deletion on the first exon and a 30-bp deletion on the third exon, respectively, for the *ms1-1* and *ms1-2* alleles [5]. An additional sample used was ‘Shindai3’ (*ms1-1*/*ms1-1*).

We applied the cetyltrimethylammonium bromide (CTAB) method, which is widely used to extract DNA from tree species. CTAB removes polysaccharides efficiently from tree tissue and recovers high quality DNA [6]. In the current study, about 100 mg of leaves, which were frozen under liquid nitrogen, were ground into fine powder with TissueLyser II (QIAGEN). The powder was washed by adding 0.9 mL of Extraction Buffer I (Additional File 1) and collected by centrifugation. The supernatant was discarded. This process was repeated if the supernatant was viscous. Then, 0.3 mL of Wash Buffer (Additional File 1) and 30 μL of 10% sodium N-dodecanoyl salcosinate were added. The mixture was left for 15 min at room temperature, before 0.3 mL of 2× CTAB buffer (Additional File 1) was added. The mixture was incubated for 10 min. at 60 °C. Then, 0.6 mL of chloroform-isoamylalchol (chloroform:isoamylalchohol=24:1) was added, aqueous layer was collected by centrifugation, and the DNA was precipitated by adding 2/3 volume of isopropanol. The precipitate was collected by centrifugation, washed by 70% ethanol, and dissolved into an appropriate volume (100 μL) of TE buffer. RNA molecules were digested by adding 1 μL of RNase (2mg/mL) and incubating for two hours at 37 °C. We used 1 μL of DNA solution (around 10—50 ng DNA) for amplification.

Because the target mutations of *MS1* gene (CJt020762, DDBJ accession numbers: LC536580 and LC538205) are deletions, we first designed PCR primers by producing products with different length by Primer3 software [7, 8]. PCR was performed in a 10 μL reaction containing 1 μL of template DNA, 1× Multiplex PCR Master Mix (Qiagen), and 0.2 μM of non-tailed primer, and 0.1 μM of each of Tail [9] and tailed primer (Table 1 and Additional File 3). The reaction was initially heat-denatured for 15 min at 95 °C, followed by 35 cycles of 94 °C for 30 sec, 60 °C for 90 sec, and 72 °C for 60 sec, with a final extension for 30 min at 60 °C, using a GeneAmp 9700 PCR System (Applied Biosystems). In addition, we synthesized specific primers to which fluorescent dye was directly attached and carried out multiplex PCR (Additional File 3) using KAPA2G Fast PCR Kit (NIPPON Genetics). The PCR products were diluted appropriately (usually by 10 times) and run on a 3130 Genetic Analyzer (Applied Biosystems) with LIZ size standard (ThermoScientific).

**Table 1.**
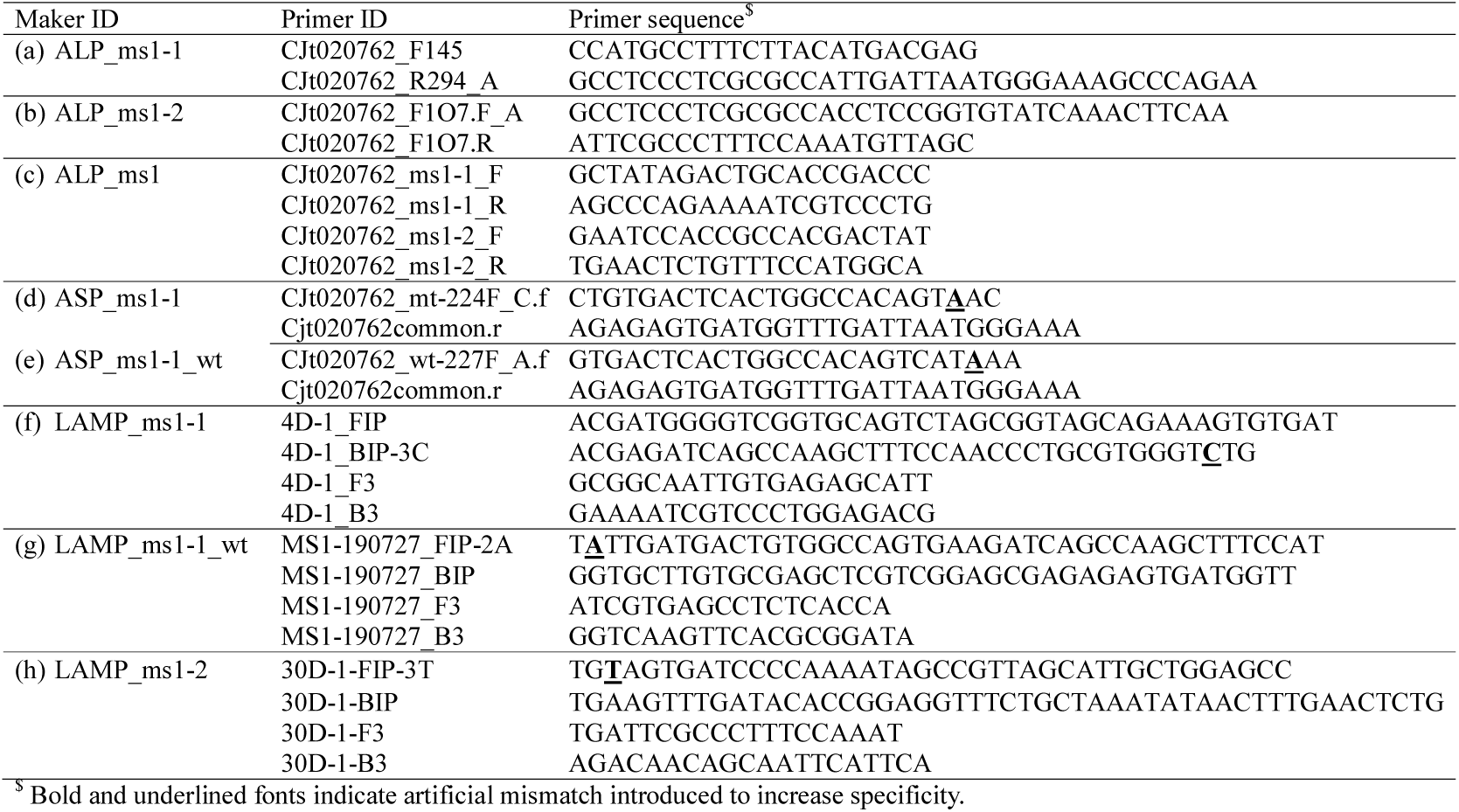
Primer sequences for PCR and LAMP diagnostic marker sets of *MS1* in *Cryptomeria japonica*.

For laboratories where capillary sequencers are not available, allele specific PCR (ASP) primers, whose products can be analyzed on agarose gels, were designed with Primer3 software [8]. One of the 3’ ends of a primer was positioned on the mismatched site due to the deletion. In addition, an artificial mismatch was introduced at the third base from 3’ end (antepenultimate position) of the primer to increase the difference in the melting temperature between matched and mismatched primer-template pair. This mismatch increased allelic specificity of the primer [10, 11]. The antepenultimate position is, for the most part, effective to discriminate different alleles. Multiplex PCR for *MS1* and one of the eight microsatellite markers in *C. japonica* (Additional File 2) were tested, and the best pair was selected, based on readability of the amplified bands on agarose gels. Because it is better to include a positive control reaction in PCR, we amplified a microsatellite marker as positive control. As a result, at least one band was visible when the product was analyzed by electrophoresis. To mitigate non-specific amplification, primers that produce shorter PCR products (with the Prime3 option: PRIMER_PRODUCT_SIZE_RANGE=‘100-250’) and shorter elongation time in PCR were selected. Reactions were carried out in 10 μL reaction containing 1 μL of template DNA, 1× Multiplex PCR Master Mix (Qiagen), and 0.2 μM of each of microsatellite forward and reverse primer, 0.2 μM of each of ASP forward and reverse primer (Table 1). The reaction was initially heat-denatured for 15 min at 95 °C, followed by 38 cycles of 94 °C for 15 sec, 63 °C for 45 sec, and 72 °C for 15 sec, using a GeneAmp 9700 PCR System (Applied Biosystems). The PCR products were separated by 2% agarose gel electrophoresis, and stained with ethidium bromide.

Primer sets for LAMP reactions were designed by PrimerExplore V5 software (https://primerexplorer.jp/e/index.html) with wild and mutant type target sequences based on ‘Fukushima1’. Design option ‘specific’ was selected, which allowed us to design allele specific LAMP primers. In order to increase the specificity of LAMP reactions and discriminate allelic types, an artificial mismatch was introduced at 3’ or 5’ position of the allele specific primer. LAMP reactions were set up in 25 μL of mixture using a DNA Amplification Kit (Eiken Chemical Co., Ltd.) according to the manufacture’s instruction. They were incubated at 63 or 65 °C for until 120 min, and deactivated at 80 °C for 5 min or 95 °C for 2 min. Fluorescent Detection Reagent (Eiken) was used to visually check the amplified products. These reactions were carried out by GeneAmp 9700 PCR System (Applied Biosystems) or real-time turbidimer, LoopampEXIA (Eiken).

## Results

The characteristics of PCR based markers are listed in Table 1. Separation of PCR products on the capillary sequencer clearly showed the length difference (in 4-bp between *ms-1* and others and 30-bp between *ms1-2* and others). The difference in length enabled the identification of the genotypes (Table 2 and Additional File 3). Moreover, for *Ms1* and *ms1-2* alleles, separation on the 2% agarose gels was attained. Products from heterozygous individual (*Ms1*/*ms1-2*) showed three bands indicating heteroduplex formation (Additional File 3).

**Table 2.**
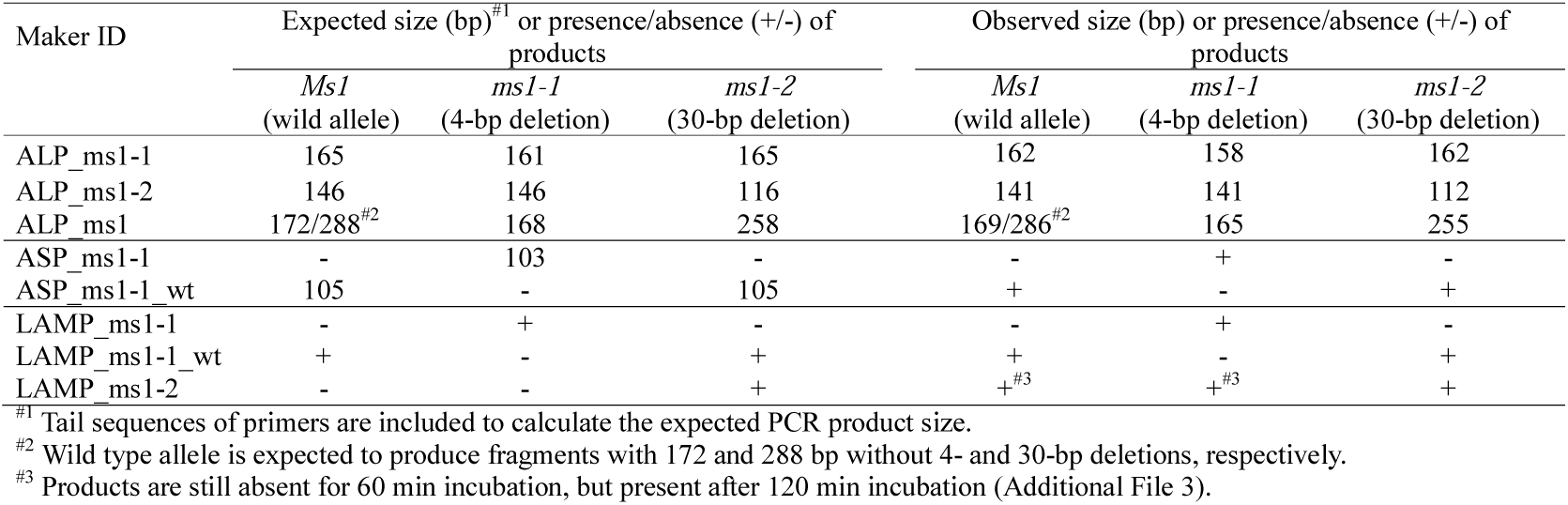
Expected and observed results of PCR and LAMP diagnostic marker for *MS1* in *Cryptomeria japonica*.

Allele specific PCR amplified the intended products allowing the discrimination of the presence or absence of the specific bands on agarose gels (Additional File 3). Under the PCR conditions described in the present study, we observed weak bands of positive control (SSR marker: CS1364) for cases where the target allele (wild or mutant allele) was present. PCR reaction mainly amplified the target with shorter PCR products. Similarly, for cases where the target alleles were absent, we observed strong single band of the positive control, this effectively suppressed the unspecific amplification from the mismatched target.

The LAMP primer sets were firstly designed with complete template-primer matching for each target allele. When possible, loop primers were also tested to speed up the LAMP reaction. Although inclusion of loop primers shortened the reaction time, they often caused false-positives. Due to these false-positives, we used primers with single artificial mismatch without loop primers. We selected the best allele specific primers (FIP: forward internal primer or BIP: backward internal primer) which showed the fastest amplification (Additional File 4).

## Discussion

We have shown that the mutant alleles for *MALE STERILITY 1* (*MS1*) can be detected in different ways and provided experimental options depending on available laboratory resources. A multiplex PCR reaction can be constructed to simultaneously detect the mutant alleles (*ms1-1* and *ms1-2*). In addition, multiplex PCR primer sets, including microsatellite markers, will be useful to manage breeding materials for male sterility. These sets enable the clonal identification of each sample. Using the allele specific primer we could locate the mutant allele among large genetic resources. Additionally, primers that amplify wild type alleles could be useful in mapping families or breeding materials, where genotyping is necessary.

In field forest stations without PCR machines LAMP reactions are convenient. As far as we know this is the first report to use LAMP for variant detection in coniferous species with a large genome (10.8 Gbp for *C. japonica* [12]). Our effort mainly focused on the detection of 4-bp deletion by LAMP because this variant is widespread in our country [5]. In addition, it is hard to separate a 4-bp difference on agarose gel electrophoresis, while 30-bp difference can be easily separated on 2% agarose gels (Additional File 3). The LAMP methods for *ms1-1* are therefore more requested than those for *ms1-2*. For LAMP, as for allele specific PCR, the use of artificial mismatch (ARMS-LAMP [13]) increased the specificity, suppressed the amplification from non-target alleles, and increased the speed of amplification for the target allele. However, complete suppression of non-target alleles was difficult (Additional File 3 (h) LAMP_ms1-2). The use of PNA (peptide nucleic acid) [14] to block the alternative allele and to increase the specificity (PNA-LAMP) may be an avenue for further study.

## Limitations

- Multiplex PCR may need optimization, depending on markers (such as microsatellites) combined and PCR machines used.
- LAMP assays may be further optimized by using other primer combinations or applying other methods, such as PNA-LAMP [14]. Sensitivity and specificity of LAMP assay has not been determined, and using both positive and negative controls is necessary to cross-validate the assay. Because we have no homozygous individuals with *ms1-2* allele and we did not construct plasmid harboring the allele, the LAMP assay to detect wild type allele without a 30-bp deletion was not reported.
- The reactions are not tested for crude DNA, for which further optimization may be needed.

## Additional files

File name: Additional File 1

File format: Excel (xlsx)

Title of data: Table S1: Buffer composition used for DNA extraction from *Cryptomeria japonica*.

Description of data: a) Extraction Buffer I, b) Wash buffer, and c) 2 x CTAB buffer.

File name: Additional File 2

File format: Excel (xlsx)

Title of data: Table S2: Microsatellite markers tested in multiplex PCR with the diagnostic ASP markers for *MS1*.

Description of data: Maker ID, Fluorescent dye, accession number, SSR motif, expected PCR product size (bp), forward primer sequence, reverse primer sequence, and reference information.

File name: Additional File 3

File format: PDF

Title of data: Figure S1 Reaction manual and representative results for diagnostic markers.

Description of data: The composition of reaction mixture, thermal conditions, and amplification results for each marker. Marker ID is the same as in Table 1 and 2.

File name: Additional File 4

File format: PDF

Title of data: Figure S2 Turbidity graph for LAMP assay with *ms1-1* specific BIP primers using Shindai3 (a: *ms1-1*/*ms1-1*) and Ooi-7 (b: *Ms1*/*ms1-2*) DNA template.

Description of data: The x-axis indicates time in minutes and the y-axis indicates turbidity.

## Abbreviations

ALP: amplified length polymorphism
ARMS: amplification refractory mutation system
ASP: allele specific PCR
BIP: backward inner primer
bp: base pair
DNA: deoxyribonucleic acid
FIP: forward inner primer
Gbp: giga base pair
LAMP: loop-mediated isothermal amplification
PCR: polymerase chain reaction
PNA: peptide nucleic acid
RNA: ribonucleic acid

## Declarations

## Availability of data and materials

The DNA sequences of CJ020762 *ms1-1* in ‘Fukushima1’, *Ms1* in ‘Ajigasawa20’ and *ms1-2* in ‘Ooi-7’ have been deposited in DDBJ with accession numbers: LC536580, LC538204, and LC538205, respectively. The datasets generated during and/or analyzed during the current study are available from the corresponding author on reasonable request.

## Authors’ contributions

YH carried out the experiments. SU carried out the experiments and wrote the manuscript. FJW identified the male sterile gene (*MS1*). AM, KU, TUI performed a part of the experiments. YM was awarded the funding, provided *Cryptomeria japonica* samples, and performed a part of the experiments. TB, KY, and SS produced genomic sequencing data for *C. japonica*. TF and MK assembled the genomic sequences. TH provided *C. japonica* samples. All authors read and approved the final manuscript.

## Acknowledgements

We thank Eiken Chemical Co., Ltd. for lending us the LoopampEXIA turbidimer, and Nozomi Ohmiya for a part of the laboratory experiments. Manuscript was checked by Crimson Interactive Pvt. Ltd. for correct grammar and spelling.

