## Additional File 3 for "Development of diagnostic PCR and LAMP markers for *MALE STERILITY 1* (*MS1*) in *Cryptomeria japonica* D. Don"

### (a) ALP_ms1-1

Amplified fragment length polymorphism (ALP) marker to detect *ms1-1* allele in *MALE STERILITY 1* (*MS1*) in sugi.

| Reaction | Volume (μL) |
|---|---|
| 2 × Multiplex (QIAGEN) | 3 |
| 2 μM CJt020762_F145 | 1 |
| 2 μM CJt020762_R294_A | 0.5 |
| 2 μM Tail_A | 0.5 |
| $dH_2O$ | 4 |
| DNA (10—50 ng) | 1 |
| Total (μL) | 10 |

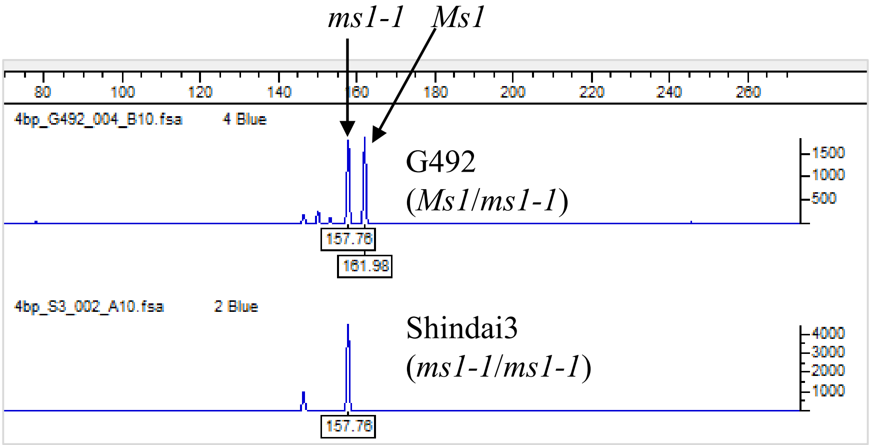


Electropherogram

| Temperature (°C) | | Cycles |
|---|---|---|
| 95 | 15 min | 1 |
| 94 | 15 sec | |
| 63 | 45 sec | 38 |
| 72 | 15 sec | |
| 60 | 30 min | 1 |
| 4 | ∞ | 1 |

| Primer ID | Primer sequence |
|---|---|
| CJt020762_F145 | CCATGCCTTTCTTACATGACGAG |
| CJt020762_R294_A | GCCTCCCTCGCGCCATTGATTAATGGGAAAGCCCAGAA |
| Tail A | [FAM]-GCCTCCCTCGCGCCA |

#### (b) ALP_ms1-2

Amplified length polymorphism (ALP) marker to detect *ms1-2* allele in *MALE STERILITY 1* (*MS1*) in sugi.

| Reaction | Volume (μL) |
|---|---|
| 2 × Multiplex (QIAGEN) | 3 |
| 2 μM CJt020762_F1O7.F_A | 1 |
| 2 μM CJt020762_F1O7.R | 0.5 |
| 2 μM Tail_A | 0.5 |
| $dH_2O$ | 4 |
| DNA (10—50 ng) | 1 |
| Total (μL) | 10 |

| Temperature (°C) | | Cycles |
|---|---|---|
| 95 | 15 min | 1 |
| 94 | 15 sec | |
| 63 | 45 sec | 38 |
| 72 | 15 sec | |
| 60 | 30 min | 1 |
| 4 | ∞ | 1 |

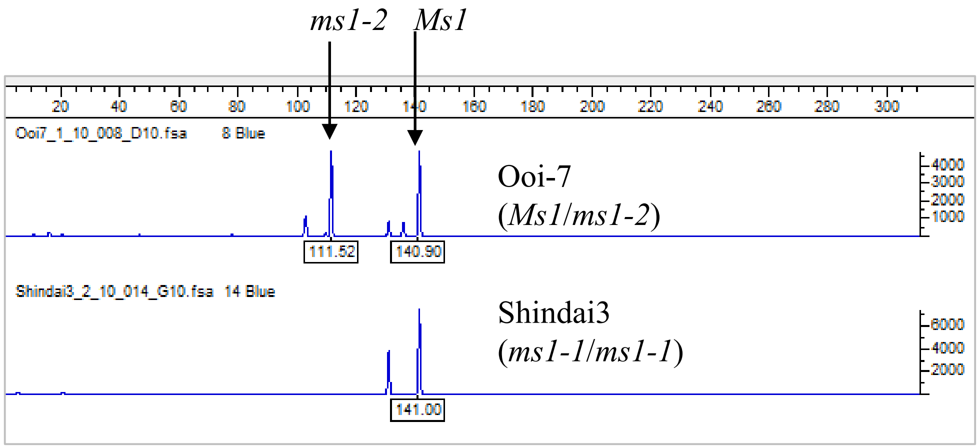


Electropherogram

| Primer ID | Primer sequence |
|---|---|
| CJt020762_F1O7.F_A | GCCTCCCTCGCGCCACCTCCGGTGTATCAAACTTCAA |
| CJt020762_F1O7.R | ATTCGCCCTTTCCAAATGTTAGC |
| Tail A | [FAM]-GCCTCCCTCGCGCCA |

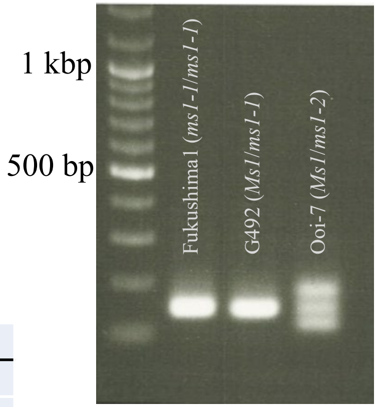


Gel image (2% agarose)

### (c) ALP_ms1

Amplified length polymorphism marker (ALP) for *MALE STERILITY 1* (*MS1*) in sugi. Two mutant alleles (*ms1-1* and *ms1-2*) and rest of the wild type alleles (*Ms1*) can be detected simultaneously by multiplex PCR.

| Reaction | Volume (mL) |
|---|---|
| KAPA2G Fast DNA Polymerase (5 U/μL) | 0.1 |
| 5 × KAPA2G Buffer A[$] | 2.0 |
| 25 mM $MgCl_2$ | 0.2 |
| dNTP Mix (10 mM each) | 0.2 |
| 5 μM CJt020762_ms1-1_F | 0.4 |
| 5 μM CJt020762_ms1-1_R | 0.4 |
| 5 μM CJt020762_ms1-2_F | 0.2 |
| 5 μM CJt020762_ms1-2_R | 0.2 |
| $dH_2O$ | 5.3 |
| DNA (5 ng/μL) | 1.0 |
| Total (mL) | 10 |

[$] 1 × KAPA2G Buffer A contains 1.5 mM $MgCl_2$

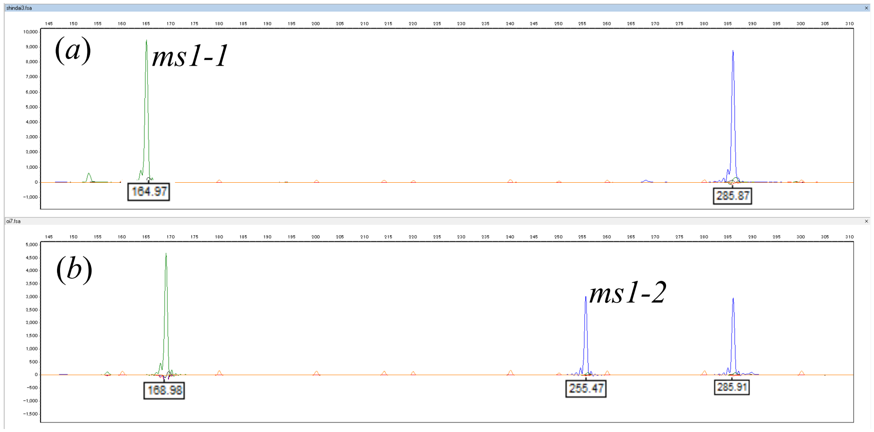


Electropherogram for (a) Shindai3 (*ms1-1*/*ms1-1*) and (b) Ooi-7 (*Ms1*/*ms1-2*)

| Temperature (°C) | | Cycles |
|---|---|---|
| 95 | 3 min | 1 |
| 95 | 15 sec | |
| 60 | 15 sec | 35 |
| 72 | 1 sec | |
| 72 | 1 min | 1 |
| 4 | ∞ | 1 |

| Primer ID | Primer sequence |
|---|---|
| CJt020762_ms1-1_F | GCTATAGACTGCACCGACCC |
| CJt020762_ms1-1_R | AGCCCAGAAAATCGTCCCTG |
| CJt020762_ms1-2_F | GAATCCACCGCCACGACTAT |
| CJt020762_ms1-2_R | TGAACTCTGTTTCCATGGCA |

**(d) ASP_ms1-1** and **(e) ASP_ms1-1_wt**

Allele specific PCR reaction to detect *ms1-1* (d) and *Ms1* (e) allele in *MALE STERILITY 1* (*MS1*) in sugi.

| Reaction | |
|---|---|
| 2 × Multiplex (QIAGEN) | 3 |
| 2 μM CJt020762_ASP$ | 1 |
| 2 μM Cjt020762common.r | 1 |
| 2 μM CS1364_F | 1 |
| 2 μM CS1364_R | 1 |
| $dH_2O$ | 2 |
| DNA | 1 |
| Total (μL) | 10 |

$ Primer ID for ASP targeting (d) *ms1-1* and (e) *Ms1* is CJt020762_mt-224F_C.f and CJt020762_WT-227F_A.f, respectively.

| Temperature (°C) | | Cycles |
|---|---|---|
| 95 | 15 min | 1 |
| 94 | 15 sec | |
| 56 | 45 sec | 38 |
| 72 | 15 sec | |
| 4 | ∞ | 1 |

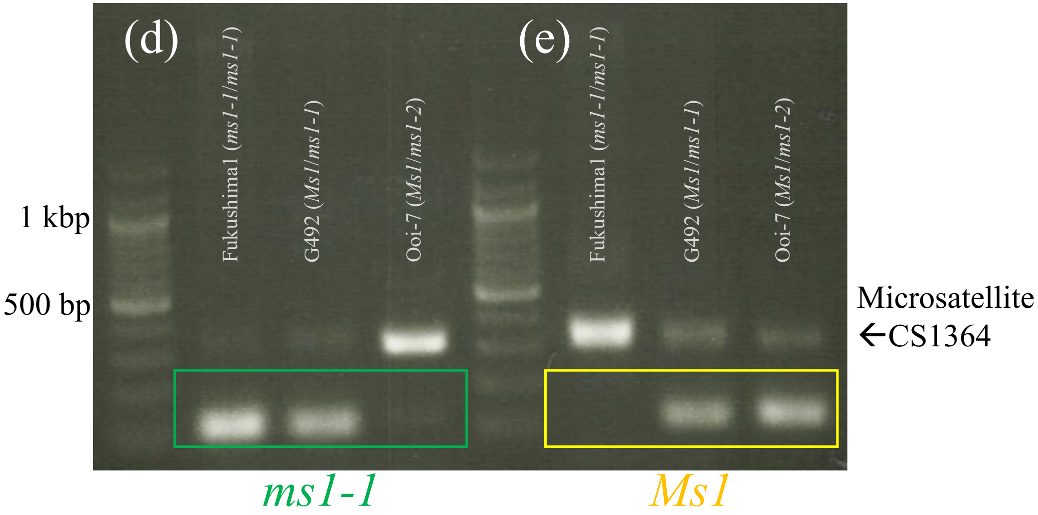


Gel image (2% agarose)

Amplification of CS1364 is suppressed if that of ASP surpasses.

| Primer ID | Primer sequence |
|---|---|
| CJt020762_mt-224F_C.f | CTGTGACTCACTGGCCACAGTAAC |
| CJt020762_WT-227F_A.f | GTGACTCACTGGCCACAGTCATAAA |
| CJt020762common.r | AGAGAGTGATGGTTTGATTAATGGGAAA |
| CS1364_F | TGATTATGGTCGGTGGTCTT |
| CS1364_R | GTGATGTGGTGTTATCTTGT |

### (f) LAMP_ms1-1

LAMP reaction to detect *ms1-1* allele in *MALE STERILITY 1* (*MS1*) in sugi.

| Reaction | |
|---|---|
| 2 x RM | 12.5 |
| 40 μM FIP (4D-1_FIP) | 1 |
| 40 μM BIP (4D-1_BIP-3C) | 1 |
| 5 μM F3 (4D-1_F3) | 1 |
| 5 μM B3 (4D-1_B3) | 1 |
| *Bst* DNA polymerase | 1 |
| DNA | 2 |
| $dH_2O$ | 5.5 |
| Total (μL) | 25 |

| Temperature ( °C) | |
|---|---|
| 63 | 120 min |
| 80 | 5 min |

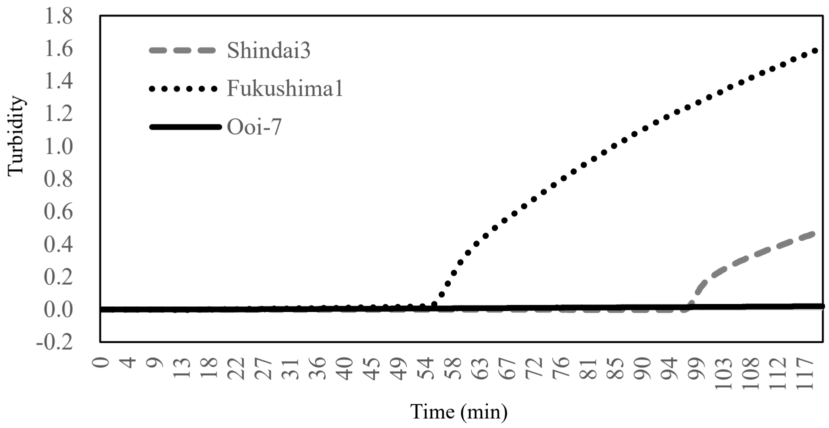


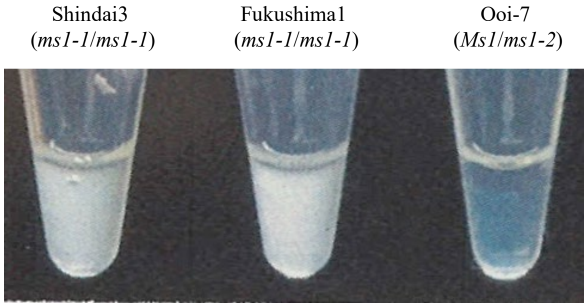


White precipitate was generated after 120 min incubation for assay with *ms1-1*.

| Primer ID | Primer sequence |
|---|---|
| 4D-1_FIP | ACGATGGGGTCGGTGCAGTCTAGCGGTAGCAGAAAGTGTGAT |
| 4D-1_BIP-3C | ACGAGATCAGCCAAGCTTTCCAACCCTGCGTGGGTCTG |
| 4D-1_F3 | GCGGCAATTGTGAGAGCATT |
| 4D-1_B3 | GAAAATCGTCCCTGGAGACG |

#### (g) LAMP_ms1-1_wt

LAMP marker to detect wild type allele (*Ms1*) without 4-bp deletion.

| Reaction | |
|---|---|
| 2xRM | 12.5 |
| 40 μM FIP (MS1-190727_FIP-2A) | 1 |
| 40 μM BIP (MS1-190727_BIP) | 1 |
| 5 μM F3 (MS1-190727_F3) | 1 |
| 5 μM B3 (MS1-190727_B3) | 1 |
| *Bst* DNA polymerase | 1 |
| DNA | 2 |
| $dH_2O$ | 5.5 |
| Total (μL) | 25 |

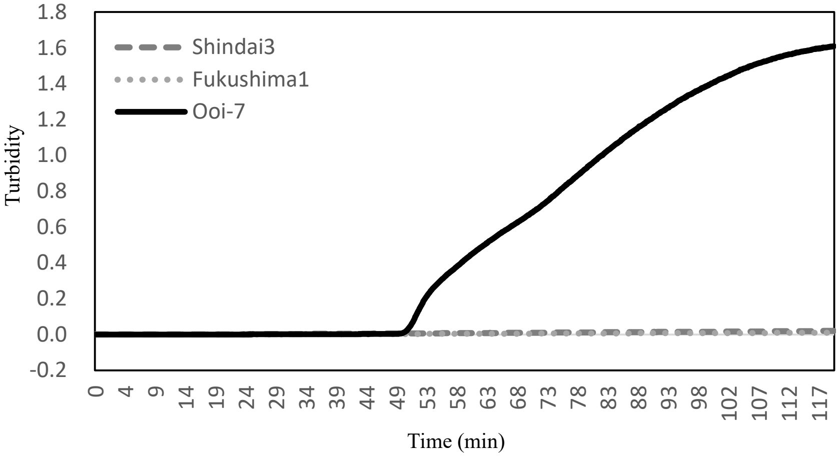


| Temperature ( °C) | |
|---|---|
| 63 | 120 min |
| 80 | 5 min |

Shindai3 (*ms1-1/ms1-1*) Fukushima1 (*ms1-1/ms1-1*) Ooi-7 (*Ms1/ms1-2*)

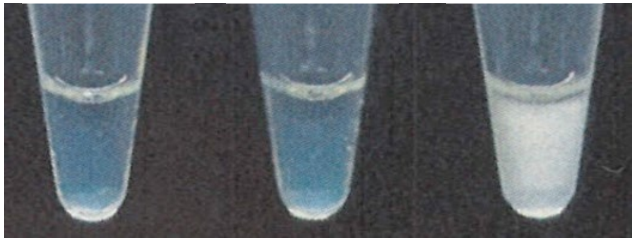

White precipitate was generated after 120 min incubation for assay with *Ms1*

| Primer ID | Primer sequence |
|---|---|
| MS1-190727_FIP-2A | TATTGATGACTGTGGCCAGTGAAGATCAGCCAAGCTTTCCAT |
| MS1-190727_BIP | GGTGCTTGTGCGAGCTCGTCGGAGCGAGAGAGTGATGGTT |
| MS1-190727_F3 | ATCGTGAGCCTCTCACCA |
| MS1-190727_B3 | GGTCAAGTTCACGCGGATA |

### (h) LAMP_ms1-2

LAMP reaction to detect *ms1-2* allele in *MALE STERILITY 1* (*MS1*) in sugi.

| Reaction | |
|---|---|
| 2xRM | 12.5 |
| 40 μM FIP (30D-1-FIP-3T) | 1 |
| 40 μM BIP (30D-1-BIP) | 1 |
| 5 μM F3 (30D-1_F3) | 1 |
| 5 μM B3 (30D-1_B3) | 1 |
| *Bst* DNA polymerase | 1 |
| DNA | 2 |
| $dH_2O$ | 5.5 |
| Total (μL) | 25 |

| Temperature ( °C) | |
|---|---|
| 65 | 120 min |
| 80 | 5 min |

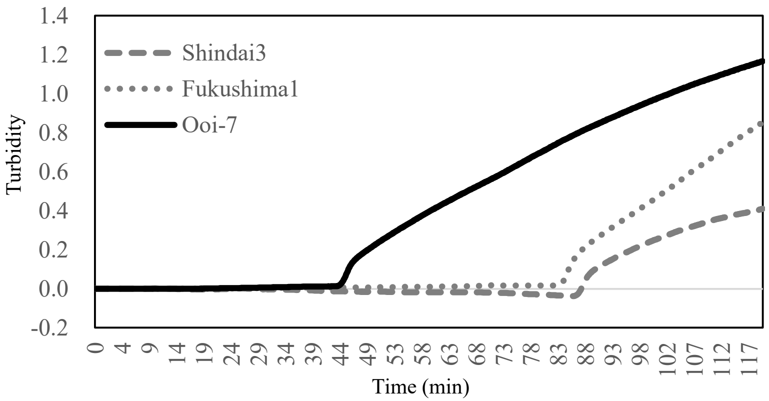


Shindai3 (*ms1-1*/*ms1-1*) Fukushima1 (*ms1-1*/*ms1-1*) Ooi-7 (*Ms1*/*ms1-2*)

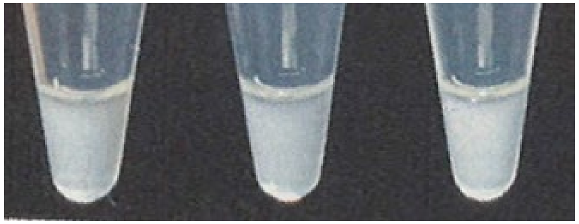

White precipitate was generated after 120 min incubation for all assays. Inclusion of positive control and stopping the reaction at 60 min. is necessary.

| Primer ID | Primer sequence |
|---|---|
| 30D-1-FIP-3T | TGTAGTGATCCCCAAAATAGCCGTTAGCATTGCTGGAGCC |
| 30D-1-BIP | TGAAGTTTGATACACCGGAGGTTTCTGCTAAATATAACTTTGAACTCTG |
| 30D-1-F3 | TGATTCGCCCTTTCCAAAT |
| 30D-1-B3 | AGACAACAGCAATTCATTCA |
