## Additional File 4 for "Development of diagnostic PCR and LAMP markers for *MALE STERILITY 1* (*MS1*) in *Cryptomeria japonica* D. Don"

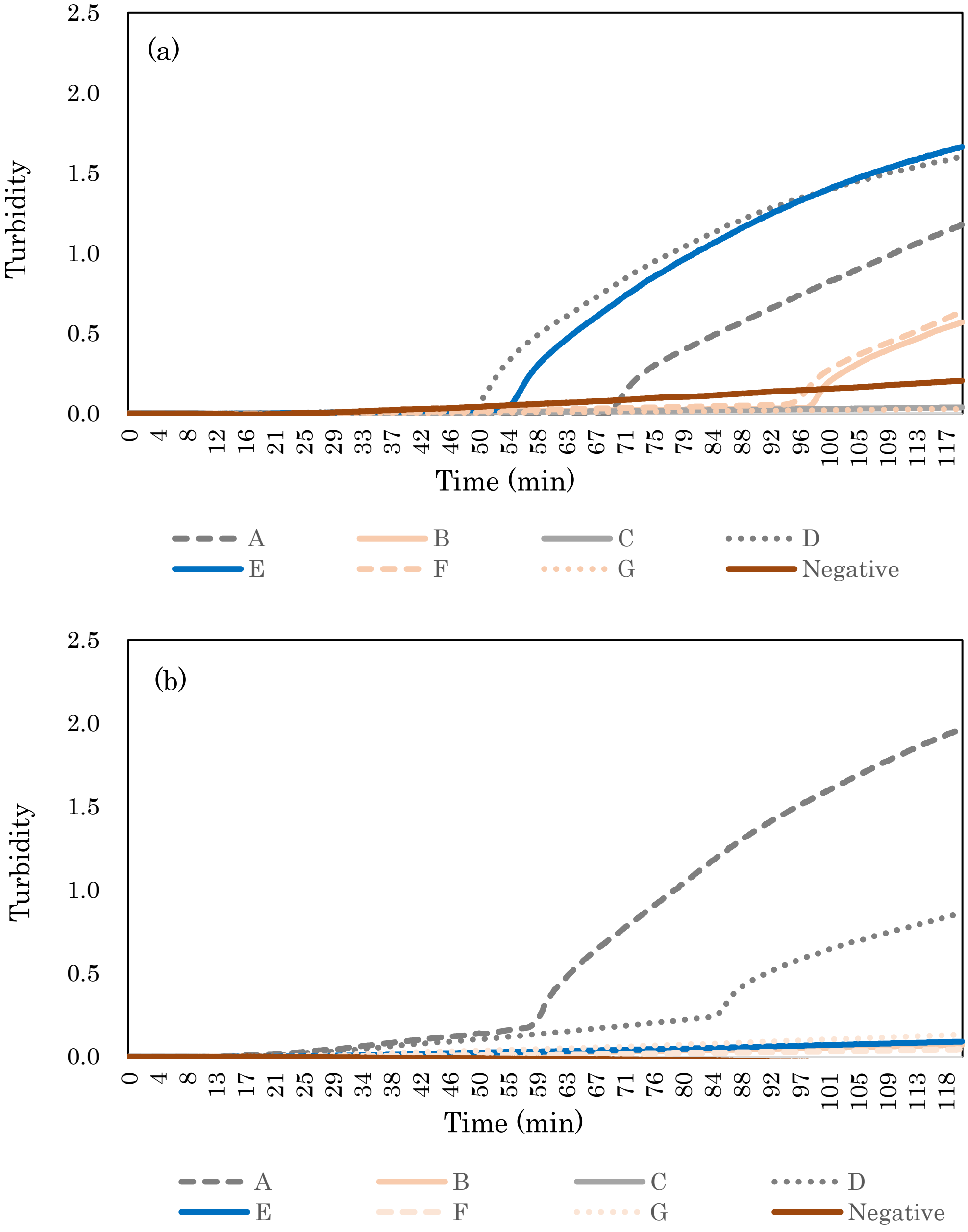


**Figure S2** Turbidity graph for LAMP assay with *ms1-1* specific BIP primers using Shindai3 (a: *ms1-1*/*ms1-1*) and Ooi-7 (b: *Ms1*/*ms1-2*) DNA template.

Each BIP primer has its own mismatched base at 3' position, with CON-XXX. CON indicates the constant part (ACGAGATCAGCCAAGCTTTCCAACCCTGCGTGGGTCTG) and XXX indicates variable bases. XXX for A to G, respectively, are GTG, GAG, GGG, GCG, CTG, ATG, TTG, where the underline indicates mismatched base introduced artificially. The BIP primer E (4D-1_BIP-3C in Table 1), shown by the blue line, shows the preferential amplification for *ms1-1*/*ms1-1* and no amplification for the *Ms1*/*ms1-2* sample. This indicates the excellent specificity of primer E for *ms1-1* compared to other primers.
